# γ-aminobutyric acid receptor B signaling drives glioblastoma in females in an immune-dependent manner

**DOI:** 10.1101/2024.07.18.603996

**Authors:** Asmita Pathak, Sravya Palasalava, Maxon V. Knott, Bruno Colon, Erika Ciervo, Yadi Zhou, Jonathan Mitchell, Oriana Teran Pumar, Harrison K. A. Wong, Li Zhang, Nikola Susic, Khushi Hemendra Shah, Kristen Kay, Diana Chin, Sadie Johnson, Feixiong Cheng, Costas A. Lyssiotis, Dionysios C. Watson, Michele Ceccarelli, Ashish Shah, Daniel R. Wahl, Justin D. Lathia, Defne Bayik

**Affiliations:** Department of Molecular & Cellular Pharmacology, Leonard M. Miller School of Medicine, University of Miami, Miami, FL, USA; Sylvester Comprehensive Cancer Center, University of Miami, Miami, FL, USA; Department of Radiation Oncology, University of Michigan, Ann Arbor, MI, USA; Leonard M. Miller School of Medicine, University of Miami, Miami, FL, USA; Università degli Studi di Napoli Federico II, Napoli, ITALY; Genomic Medicine Institute, Lerner Research Institute, Cleveland Clinic, Cleveland, OH, USA; Department of Molecular & Integrative Physiology, University of Michigan, Ann Arbor, MI, USA; Rogel Cancer Center, University of Michigan, Ann Arbor, MI, USA; Department of Cardiovascular & Metabolic Sciences, Lerner Research Institute, Cleveland Clinic, Cleveland, OH, USA; Case Comprehensive Cancer Center, Cleveland, OH, USA; Department of Molecular Oncology, Leonard M. Miller School of Medicine, University of Miami, Miami, FL; Department of Neurosurgery, Leonard M. Miller School of Medicine, University of Miami, Miami, FL; Rose Ella Burkhardt Brain Tumor & Neuro-Oncology Center, Cleveland Clinic, Cleveland, OH, USA

**Keywords:** glioblastoma, GABA, sex differences, myeloid-derived suppressor cells, L-arginine, nitric oxide synthase 2

## Abstract

Sex differences in immune responses impact cancer outcomes and treatment response, including in glioblastoma (GBM). However, host factors underlying sex specific immune-cancer interactions are poorly understood. Here, we identify the neurotransmitter γ-aminobutyric acid (GABA) as a driver of GBM-promoting immune response in females. We demonstrated that GABA receptor B (GABBR) signaling enhances L-Arginine metabolism and nitric oxide synthase 2 (NOS2) expression in female granulocytic myeloid-derived suppressor cells (gMDSCs). GABBR agonist and GABA analog promoted GBM growth in females in an immune-dependent manner, while GABBR inhibition reduces gMDSC NOS2 production and extends survival only in females. Furthermore, female GBM patients have enriched GABA transcriptional signatures compared to males, and the use of GABA analogs in GBM patients is associated with worse short-term outcomes only in females. Collectively, these results highlight that GABA modulates anti-tumor immune response in a sex-specific manner, supporting future assessment of GABA pathway inhibitors as part of immunotherapy approaches.

## INTRODUCTION

Glioblastoma (GBM) is the most common primary malignant brain tumor in adults, with a median survival of ∼18-20 months^1^. Despite promising advances in solid tumor therapy with immune checkpoint inhibitors, GBM remains highly resistant to these treatment strategies^2–6^. This arises in part from the immunosuppressive GBM microenvironment that is abundant in myeloid cell populations^7,8^. Among them, myeloid-derived suppressor cells (MDSCs) are a heterogeneous population of bone marrow-derived cells consisting of monocytic (mMDSC) and granulocytic (gMDSC) subsets, which differ in their suppressive activity and tumor interactions^9–12^. Increased peripheral circulation and tumor infiltration of MDSCs correlate with poor GBM prognosis^13–15^ highlighting the importance of MDSC-mediated immune suppression in this disease. However, limited knowledge of the factors driving MDSC accumulation and function prevents effective targeting of these cells in GBM. There is growing evidence that sex differences in immune response further inform GBM outcomes and potential treatment targets^10,16–18^. We previously demonstrated that MDSC subsets play sex-specific roles in GBM since gMDSC expansion drives GBM in female preclinical models and a high gMDSC signature correlates with worse prognosis in female patients^10^. However, host mechanisms informing the distinct function of MDSCs in GBM remain poorly investigated.

Beyond immune interactions, the communication between GBM cells and the neuronal environment impacts tumorigenesis. In preclinical models of brain tumors and GBM patients, neurons were shown to drive GBM growth and invasion through direct electrochemical interactions or via secretion of neurotransmitters and growth factors^19–25^. However, the role of neurotransmitters can be contextual. Aberrant γ-aminobutyric acid (GABA) signaling is one such example, as GABA receptor A (GABA_A_R) was shown to suppress the proliferation of GBM cells while supporting the metastasis of breast cancer to the brain^26–29^. In addition to these direct cancer cell effects, emerging studies indicate that GABA can modulate anti-tumor immunity. Elevated production of GABA from cancer cells or B cells induces immunosuppressive macrophage polarization and/or suppresses T cells in colon and lung cancers^30,31^. However, the role of GABA in GBM immune response remains unknown. Here, we demonstrate that GABA drives GBM in a sex-specific manner by metabolically programming gMDSCs in females. Our findings identify GABA receptor B (GABBR) signaling as a therapeutically relevant pathway in females.

## RESULTS

### gMDSCs express GABA receptors

While our earlier studies demonstrated that mMDSCs and gMDSCs play a differential role in GBM^10^, with gMDSCs having an enrichment in females, the pathways contributing to their distinct function remain to be investigated. Recent advances identified targetable pathways in mMDSCs to enhance immunotherapy efficacy^32–34^, while strategies to target gMDSCs in GBM are lacking. Thus, to identify mechanisms that are linked to gMDSC functionality, we re-analyzed the list of drug candidates predicted to target individual MDSC subsets based on their differential expression signatures (GSE148467) ^10^. Among the top 20 clinically used drugs that were predicted to be selective for gMDSCs, three were identified as GABA pathway modulators (**Fig.1a, Extended Fig. 1a**). The other top-ranked drug classes included G-CSF and IL-1β modulators, which have been previously shown to be associated with gMDSC activity and identified as potential immunotherapy targets^10,35,36^ Therefore, we evaluated whether MDSCs are capable of responding to GABA by analyzing the expression levels of GABA_A_R, an ion channel, versus GABBR, a G protein-coupled receptor^37^. We observed that both GABBR heterodimer subunits GABBR1, and GABBR2, and GABA_A_R2 expression was higher in mouse bone marrow-derived gMDSCs compared to mMDSCs (**Fig.1b-c**). Since our previous results demonstrated that gMDSC signature in tumors correlates with worse outcomes specifically in female GBM patients^10^, we analyzed sex differences in GABA signaling transcriptional profiles using a publicly-available single-cell RNA-sequencing dataset (GSE117891)^38^. We observed that tumor-infiltrating immune cells in female patients had enriched GABA signature along with higher GABBR and GABA_A_R expression compared to male patients (**Fig. 1d, Extended Fig. 1b**). Notably, the mean expression of GABBR was higher than that of GABA_A_R (**Extended Fig. 1b).** Therefore, we analyzed differential GABBR expression across immune lineages. Tumor-associated macrophages, microglia, and neutrophils/gMDSCs had significantly higher GABBR1 and GABBR2 expression in females compared to males, while there was no sex difference in the expression levels of GABBR in monocytes/mMDSCs (**Fig. 1e-f**). Collectively, our data suggest that GABA could be a potential modulator of gMDSC activity.

**Figure 1:**
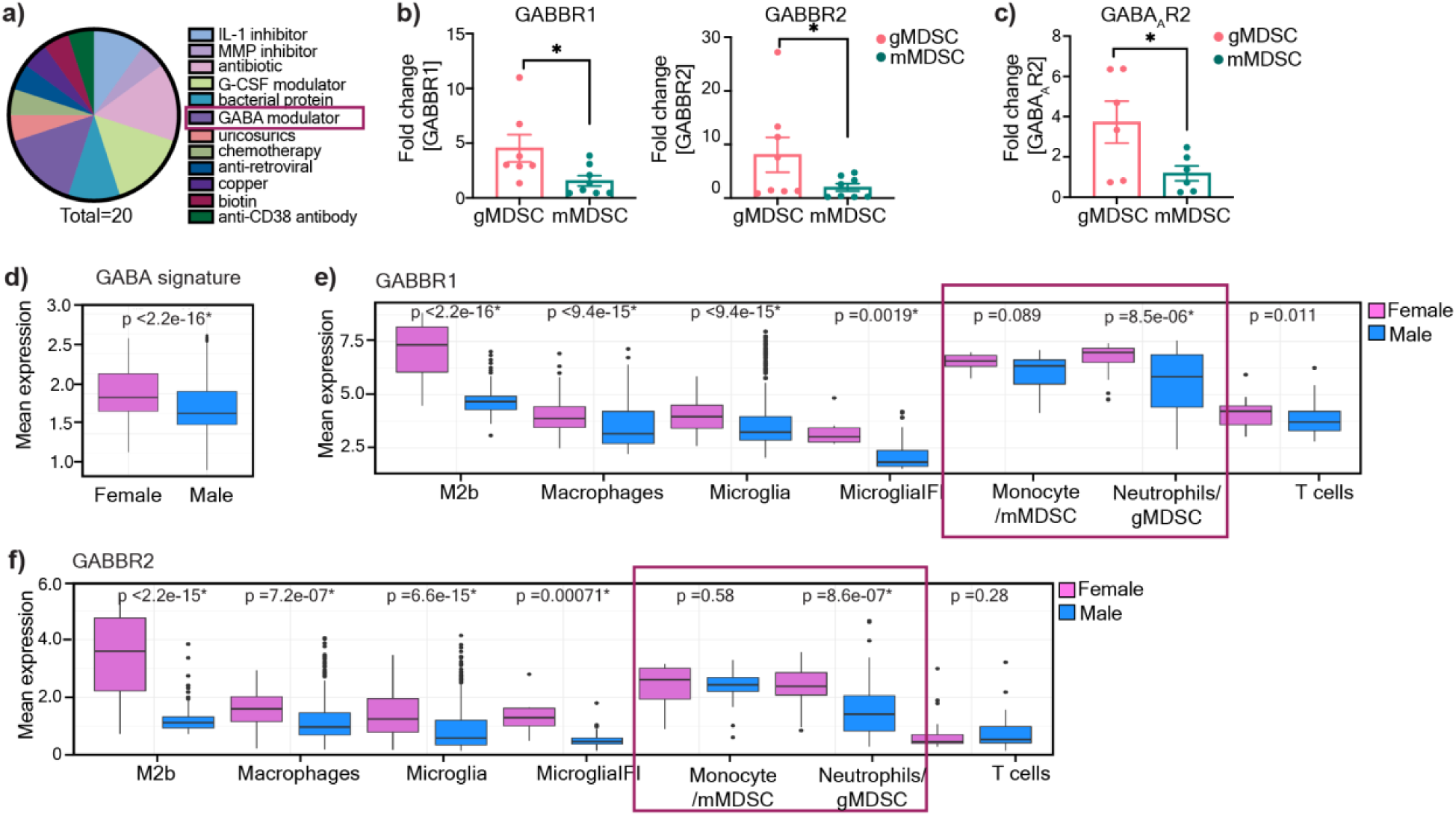
gMDSCs express GABA receptors. **a)** Pie chart illustrating the distribution of molecular targets among 20 clinically-approved drug candidates predicted to be effective against gMDSCs. **b)** Graph showing the expression levels of GABBR1 (left) and GABBR2 (right) in mouse bone marrow-derived gMDSCs and mMDSCs (n=7-8; *p < 0.05 as determined by unpaired t-test) **c)** Expression of GABA_A_R2 in mouse bone marrow-derived gMDSCs and mMDSCs (n=4). Data shown as mean ± SEM. **d)** GABA signature (GABA_A_R, GABBR, GABA transaminase, and GABA transporter in tumor infiltrating immune populations (macrophages, microglia, monocytes/mMDSCs, neutrophils/gMDSCs and T cells) in female (n=5) and male (n=8) glioma patients along with **e)** Mean expression levels of GABBR1 (top panel) and GABBR2 (bottom panel) in female (n=5) and male (n=8) immune populations of glioma patients. *p<0.05 as determined by Wilcoxon Rank-Sum test.

### GABA signaling enhances L-arginine metabolism in female gMDSCs

Based on the differential receptor expression profile in MDSCs and previous studies that demonstrated GABA signaling alters immune cell metabolism^31,39^ we tested whether GABA metabolically reprograms gMDSC. A targeted metabolite screen demonstrated that overnight treatment of MDSC subsets with GABA led to intracellular accumulation of L-Arginine in female gMDSCs (p-adj<0.01), with no notable metabolic alterations following GABA stimulation in any of the other MDSC subsets (**Fig. 2a, Extended Fig. 2a**). L-Arginine is an essential amino acid that is critical for the immunosuppressive function of MDSCs^40^. Importantly, untreated female gMDSCs had significantly lower levels of L-Arginine compared to male counterparts, consistent with lower baseline expression of the L-Arginine transporter Cat2b^41^, which was upregulated in female gMDSCs upon GABA stimulation (**Extended Fig. 2b-d**). Correspondingly, L-arginine biosynthesis and arginine metabolism emerged as the top upregulated metabolic pathways in GABA-treated female gMDSCs (**Fig. 2b**). Importantly, there was a similar increase in GABA levels across mMDSCs and gMDSCs from either sex, suggesting that the effect is likely mediated through signaling rather than differences in intracellular accumulation (**Extended Fig. 2e**).

**Figure 2:**
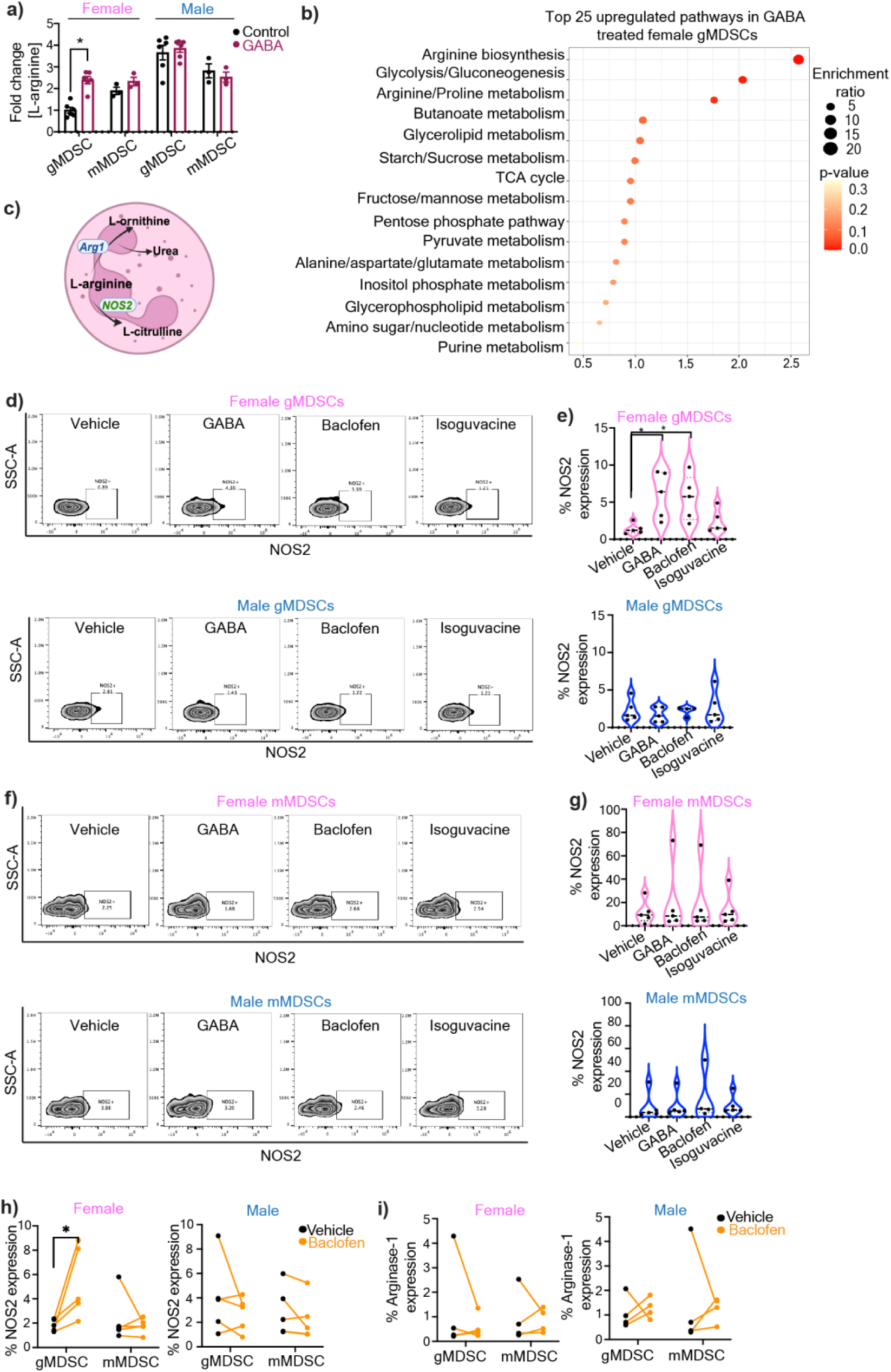
GABA metabolically alters female gMDSCs through upregulation of L-arginine. **a)** gMDSCs (CD11b^+^Ly6G^+^Ly6C^-^) and mMDSCs (CD11b^+^Ly6G^-^Ly6C^+^) from male and female mouse were sorted from bone marrows and treated with 100 μM GABA overnight. Relative L-Arginine levels in female and male gMDSCs (n=6) and mMDSCs (n=3) following GABA treatment were determined by mass spectrometry. *p<0.01 as determined by 2-way ANOVA **b)** Graphical representation depicting the metabolic conversion of L-arginine to L-ornithine and L-citrulline by Arginase-1 (Arg-1) and nitric oxide synthase 2 (NOS2), respectively. Male and female mouse MDSCs were treated with 100 μM GABA, baclofen, isoguvacine or PBS (vehicle) overnight. **c)** NOS2 expression was analyzed by flow cytometry. Representative contour plots demonstrating the change in NOS2 expression in gMDSCs with GABA, baclofen, and isoguvacine treatment. **d)** Violin plots showing the percentage of NOS2 expression in female (pink, top) and male (blue, bottom) mouse gMDSCs (n=4-5). *p< 0.05 as determined by unpaired t-test**. e)** Representative contour plots demonstrating the change in NOS2 expression in mMDSCs with GABA, baclofen, and isoguvacine treatment. **f)** Violin plots depicting the percentage of NOS2 expression in female (pink, top) and male (blue, bottom) mouse mMDSCs (n=4-5). **g)** Representative contour plots demonstrating the change in Arg1 expression in gMDSCs with GABA, baclofen, and isoguvacine treatment. **h)** Violin plots showing the percentage of Arg1 expression in female (pink, top) and male (blue, bottom) mouse gMDSCs (n=4). **i)** Representative contour plots demonstrating the change in Arg1 expression in mMDSCs with GABA, baclofen, and isoguvacine treatment. **j)** Violin plots depicting the percentage of Arg1 expression in female (pink, top) and male (blue, bottom) mouse mMDSCs (n=4). Human PBMCs were treated with 100 μM baclofen or PBS (vehicle) overnight and expression of NOS2 and Arg-1 analyzed by flow cytometry. The frequency of **k)** NOS2 (n=5) and **l)** Arg-1 (n=4) expressing gMDSCs versus mMDSCs from female (left) and male (right) donors. *p<0.05 as determined by paired t-test.

L-Arginine is metabolized by either Arginase-1 (Arg-1) or nitric oxide synthase 2 (NOS2) to yield L-ornithine or L-citrulline/nitric oxide (NO), respectively^40–42^ (**Fig. 2c**). Since GABA treatment of gMDSCs increased L-citrulline (p<0.05), and NO was shown to enhance GBM tumorigenicity^43,44^, we initially investigated NOS2 as a potential mediator of GABA signaling (**Extended Fig. 2a**). There was a significant increase in NOS2 protein levels in female mouse gMDSCs treated with GABA or the GABBR agonist baclofen, but not with the GABA_A_R agonist isoguvacine (**Fig. 2d-e**). In contrast, we did not observe any discernible differences in NOS2 expression of male gMDSCs or mMDSCs of either sex **(Fig. 2d-g**). Similarly, Arg-1 expression remained unchanged in both male and female MDSC subsets stimulated with GABA or GABA receptor agonists (**Extended Fig. 2f-i**). To validate whether GABA-mediated NOS2 upregulation is conserved in humans, we tested the effect of GABBR signaling in MDSCs from human peripheral blood mononuclear cells (PBMC). Consistent with our mouse MDSC data, baclofen stimulation induced NOS2 but not Arg-1 expression specifically in female gMDSCs from healthy donors (**Fig. 2h-i, Sup. Fig 1**). Overall, these results indicate that GABBR signaling in female gMDSCs enhances the activity of the NOS2 pathway.

### GABBR drives GBM in females

MDSCs mediate their immunosuppressive activity by suppressing T cell proliferation through NO-dependent mechanism^45^. Therefore, we examined the impact of GABBR signaling on MDSC-mediated T cell proliferation. Baclofen treatment alone had no effect on the proliferation rate of T cells directly nor did it increase mMDSC-mediated T cell suppression (**Fig. 3a-b**). However, female gMDSCs treated with baclofen acquired enhanced T cell suppressive properties compared to vehicle controls (**Fig. 3a**). This effect was dependent on NOS2, as its inhibition by aminoguanidine reversed baclofen-induced T cell suppression of female gMDSCs (**Fig. 3c**). Importantly, consistent with the existing literature demonstrating the role of NOS2 in mMDSC-mediated suppression^45^, aminoguanidine treatment significantly reduced the suppressive capacity of mMDSCs (**Fig. 3c-d**). Since we also observed higher mean GABBR expression in female macrophages in addition to granulocytes (**Fig. 1e-f**) and previous reports suggested that GABA could impact macrophages in colon cancer^31^, we also analyzed the potential effect of GABA on mouse macrophage functionality by interrogating the expression levels of canonical markers associated with pro-inflammatory (I-A/I-E, CD80, CD86) or immunosuppressive (CD204, CD206) phenotypes. There was no discernible difference in expression levels of any of these markers with GABA treatment (**Extended Fig. 3a**).

**Figure 3:**
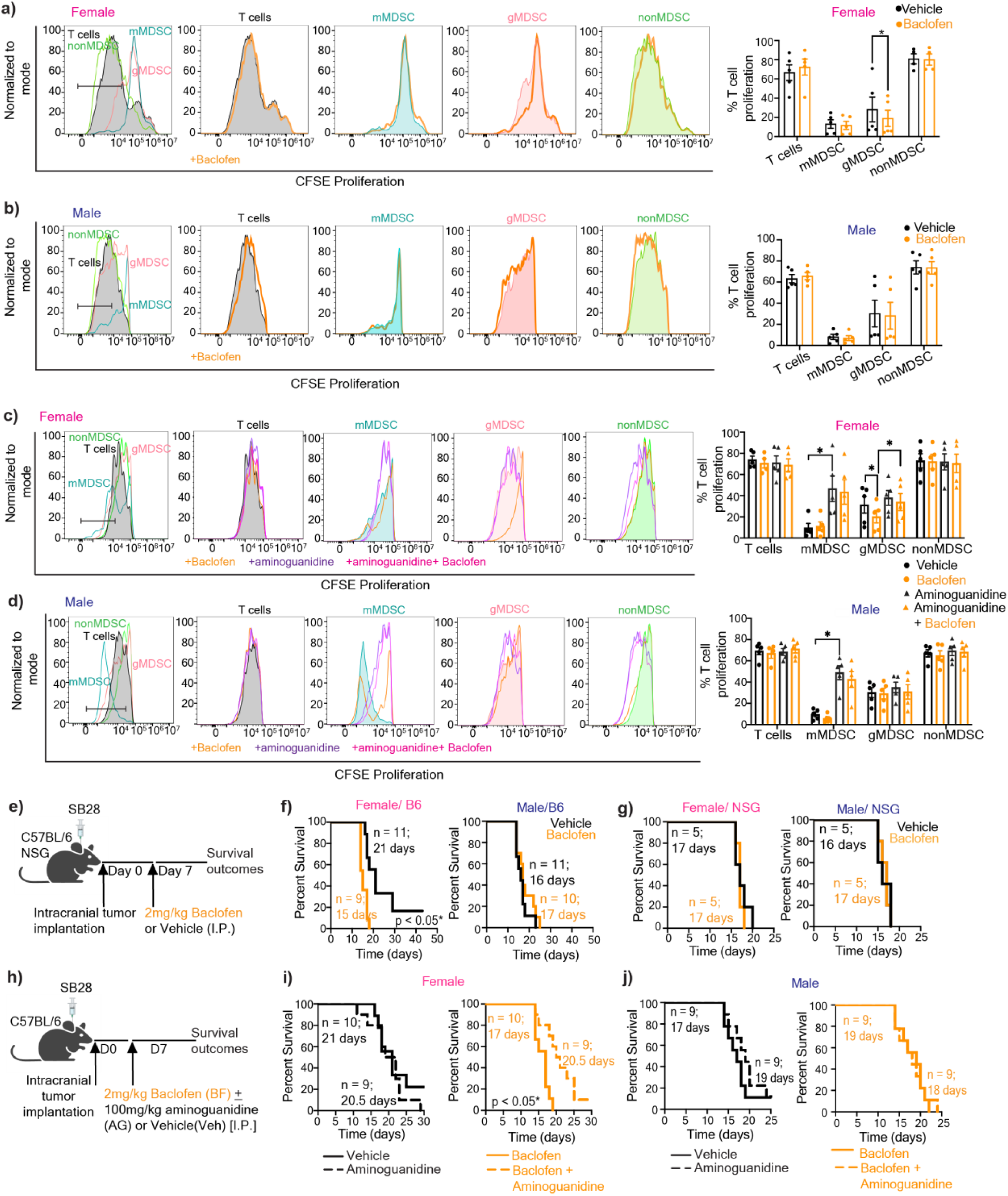
GABBR signaling drives GBM and gMDSC-mediated T cell suppression in females in a NOS2-dependent manner. T cells activated with 100IU/ml IL-2 and anti-CD3/CD28 were co-cultured with mMDSCs (CD11b^+^Ly6G^-^Ly6C^+^), gMDSCs (CD11b^+^Ly6G^+^Ly6C^-^), nonMDSCs (CD11b^+^Ly6G^-^Ly6C^-^). T cells and myeloid co-cultures were treated daily with 100 μM baclofen. Representative histograms depicting proliferation rate of T cells (left), and graphs showing the frequency of proliferating T cells (right) for **a)** female, and **b)** male samples. Mean + SEM combined from n=4-5 samples; *p<0.05 as determined by ratio paired t-test. T cells and myeloid cell cocultures were treated with baclofen daily and/or 1mM aminoguanidine once. Representative histograms depicting proliferation rate of T cells (left), and graphs showing the frequency of proliferating T cells (right) for **c**) females and **d)** males, n=5, *p<0.05 as determine by paired t-test. **e**) Female and male C57BL/6 mice were intracranially injected with 30,000 (females) and 20,000 (males) SB28 cells. NSG mice were orthotopically implanted with 10,000 SB28 cells. 7 days post-tumor implantation mice were intraperitoneally injected with 2 mg/kg baclofen or vehicle following a 5-days-on-2-day-off schedule. Kaplan-Meier curves depicting median survival span of female (left) and male (right) **f)** C57BL/6 mice (data combined from two independent experiments), and **g)** NSG mice treated with vehicle versus baclofen. **h)** Female and male C57BL/6 mice were intracranially injected with 30,000 (females) and 20,000 (males) SB28 cells. 7 days post-tumor implantation mice were intraperitoneally injected with 2 mg/kg baclofen and/or 100 mg/kg aminoguanidine following a 5-days-on-2-day-off schedule. Kaplan-Meier curves depicting median survival span of **i)** female and **j)** male mice treated with aminoguanidine alone (left) or baclofen plus aminoguanidine (right). Data were combined from two independent experiments. * p<0.05 based on the log-rank test.

Since GABA was shown to interfere with anti-tumor T cell response^30,31^ we evaluated whether GABBR signaling impacts tumorigenesis in preclinical models of GBM. Female immunocompetent mice orthotopically implanted with the syngeneic GBM model SB28 and treated with baclofen succumbed to GBM earlier compared to vehicle-treated mice (**Fig. 3e-f**). This intervention did not impact disease outcomes in immunocompetent male mice or immunocompromised mice of either sex, highlighting the role of the intact immune system in baclofen-mediated tumorigenesis (**Fig. 3f-g**). GABA or baclofen also had no effect on the proliferation rate of syngeneic mouse GBM lines or patient-derived xenografts (PDXs) *in vitro* (**Extended Fig. 3b**). Considering the increased NOS2 expression in female gMDSCs with GABA and baclofen treatment, we sought to determine whether NOS2 acts as a downstream effector of baclofen *in vivo*. Inhibition of NOS2 by aminoguanidine abrogated the protumorigenic effect of baclofen in female mice with GBM, while it had no effect on survival of male mice or vehicle - treated female mice (**Fig. 3h-j**). Taken together, these results suggest that GABBR signaling enhances gMDSC-mediated immunosuppression in females and accelerates GBM in a NOS2-dependent manner.

### GABA analog use associates with worse outcomes in female GBM patients

Since GABA analogs are used in cancer patients to manage neuropathic pain^46^, we tested the effect of one of the GABA analogs, pregabalin, in syngeneic mouse models. We observed that pregabalin treatment significantly reduced median survival duration in immunocompetent female mice bearing SB28 or KR158 tumors (**Extended Fig. 3c-d**). In contrast, pregabalin had no effect on the survival span of male immunocompetent mice or immunocompromised mice of either sex (**Extended Fig. 3e-f**). To determine the clinical relevance of these observations, we used a case-control type study design and interrogated whether commonly used GABA analogs (gabapentin and pregabalin) impact GBM patient survival after diagnosis. We used date of initial surgical resection (standard of care upon presentation of GBM) and date of death to determine overall survival duration. Among 990 brain tumor patients in our institutional registry, we identified 169 patients with a history of either pregabalin or gabapentin use. After applying inclusion and exclusion criteria detailed in the Methods section, we assembled a retrospective cohort of primary GBM patients consisting of 18 males and 23 females on GABA analogs, and 111 males and 94 females not on these drugs. We observed that females on drugs had significantly lower 12-month overall survival compared to those not on drugs (0.41 vs. 0.65, p=0.029) (**Fig. 4a**). There was no effect of GABA analog treatment on 12-month overall survival in males (0.65 vs. 0.63, p=0.946) (**Fig. 4b**).

**Figure 4:**
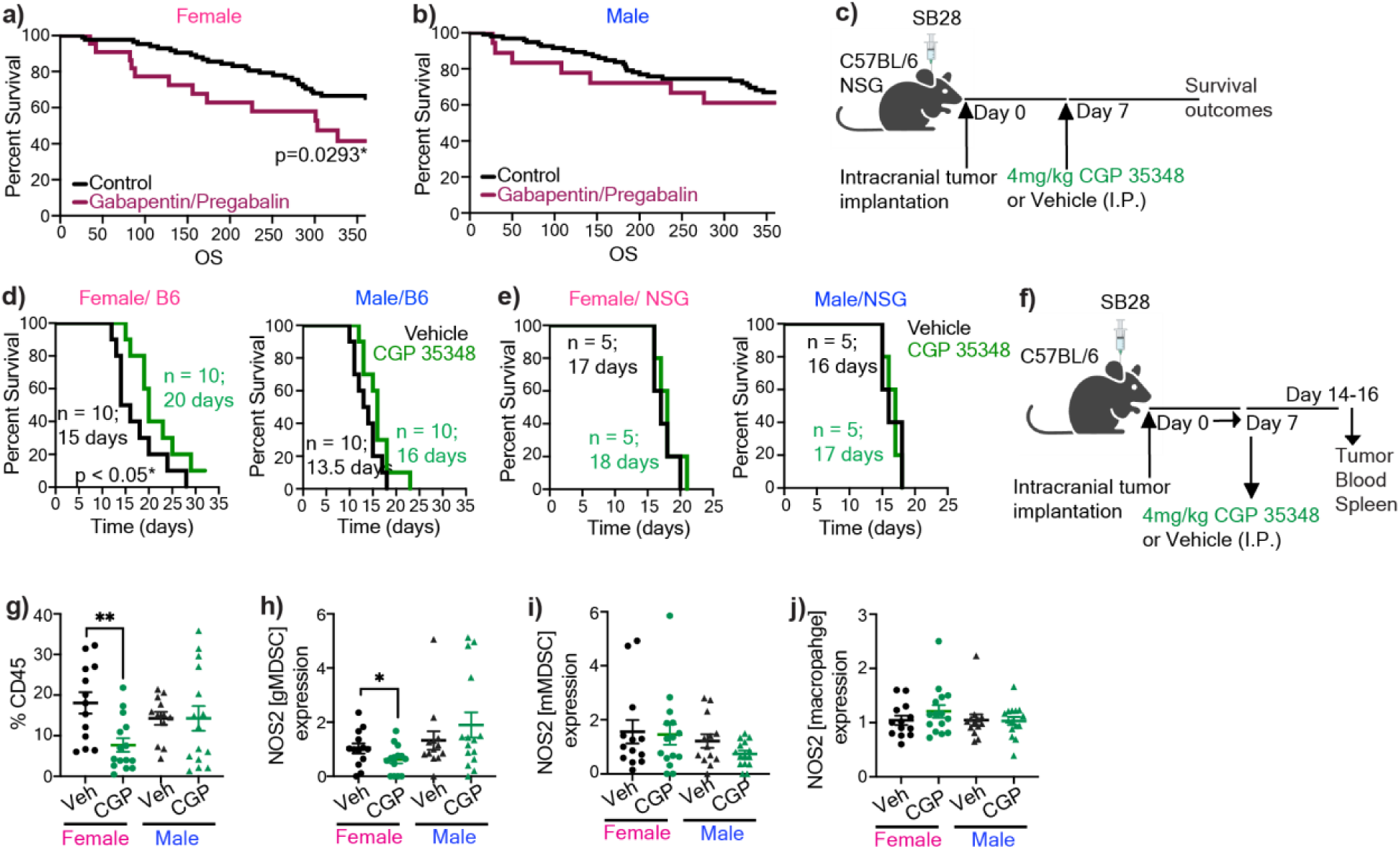
Modulation of GABBR signaling impacts GBM survival in females highlighting its clinical relevance. Kaplan-Meier curves depicting survival of **a)** female and **b)** male patients with primary GBM on gabapentin/pregabalin versus the control cohort. p<0.05 as determined by log-rank test. **c)** Female and male C57BL/6 mice were intracranially implanted with 30,000 (females) and 20,000 (males) SB28. NSG mice were implanted with 10,000 SB28. 7 days post-tumor implantation mice were intraperitoneally injected 4 mg/kg CGP 35348 or vehicle following a 5-days-on-2-day-off schedule. Kaplan-Meier curves depicting median survival span of female (left) and male (right) **d)** C57BL/6 mice (data were combined from two independent experiments), and **e)** NSG mice. * p<0.05 as determined by log-rank test. **f)** Schematics illustrating the timeline for the analysis of the immune landscape following CGP 35348 treatment of SB28-bearing male and female mice. **g)** The percentage of tumor-infiltrating CD45+ cells, and the relative frequency of **h)** NOS2+ gMDSCs from tumors, **i)** NOS2+ mMDSCs from tumors, **j)** NOS2+ macrophages from tumors of vehicle or CGP 35348 treated mice (n=13-15). Data shown from individual animals, combined from three independent experiments. *p<0.05 determined by unpaired t-test.

### GABBR as an immunotherapy target

As our results highlighted that the GABA-GABBR axis promotes GBM in females, we hypothesized that inhibiting GABBR signaling could serve as a potential therapeutic intervention. We systemically treated SB28-bearing mice with the GABBR antagonist CGP 35348 (**Fig. 4c**). This strategy significantly improved survival in immunocompetent female mice, whereas it did not affect survival of males. We also did not observe an effect on survival in immunocompromised mice, consistent with an immune-dependent effect of CGP 35348 treatment in GBM (**Fig. 4d-e**). This prompted us to evaluate the immune landscape of CGP 35348 versus vehicle-treated animals (**Fig. 4f**). Administration of CGP 35348 led to a significant reduction in overall immune cell infiltration in female tumors (**Fig. 4g**). While there was no significant change in the relative proportion of gMDSCs or any other immune population, there was a significant reduction in NOS2 expression from tumor-infiltrating gMDSCs in females (**Fig. 4h**, **Extended Fig. 4**). This NOS2 effect was not observed in tumor-infiltrating mMDSCs or macrophages (**Fig. 4i-j**). Moreover, there were no differences in the frequency of peripheral immune populations or NOS2 production from circulating or splenic gMDSCs (**Extended Fig.5**, **Sup Fig. 2**). Collectively, these data support the conclusion that GABBR inhibition improves survival in females with GBM by reducing NOS2 production from tumor-infiltrating gMDSCs (**Fig. 5**).

**Figure 5:**
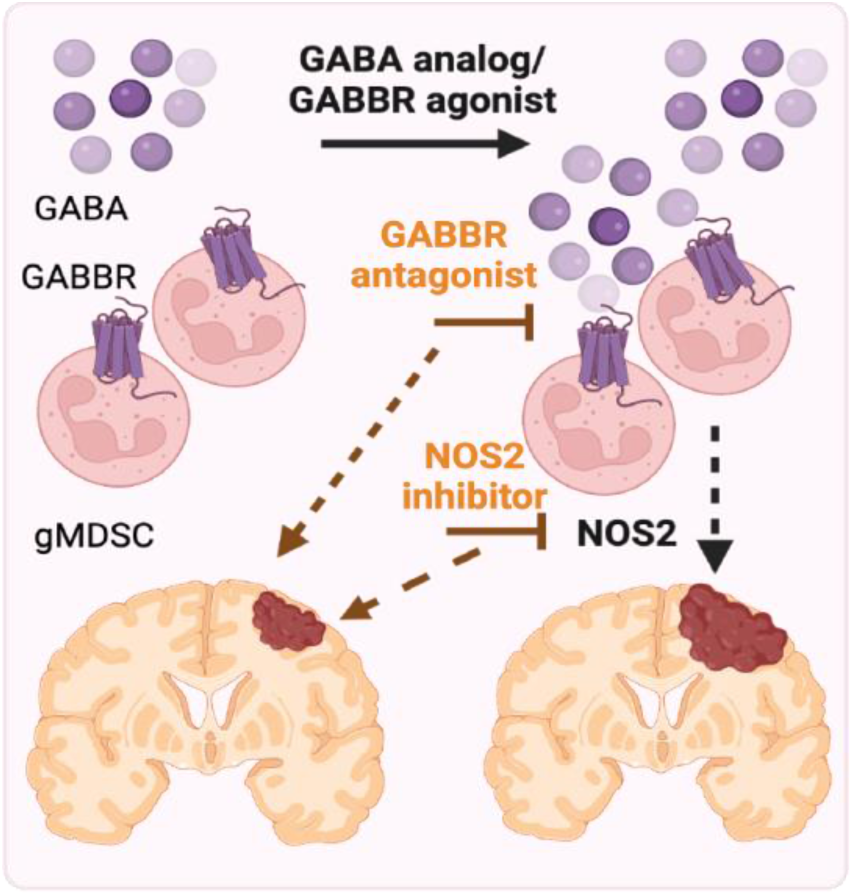
GABA-GABBR axis promotes NOS2-mediated GBM in females. Proposed model depicting the effect of GABA in the female GBM microenvironment. Model created with biorender.

## DISCUSSION

The communication between the nervous system and cancer cells in GBM has revealed a previously underappreciated aspect of tumor biology^19–21,24,25^. This effect extends beyond brain tumors as neurons directly impact tumor growth and metastasis through synaptic-like interactions and neurotransmitter release in various solid tumors^47–50^. However, the mechanisms by which neuronal signaling reshape the non-malignant components of the tumor microenvironment remain largely unexplored. In this study, we observed that GABA regulates anti-GBM immune responses in a sex-specific manner by enhancing the immunosuppressive function of gMDSC through upregulation of the NOS2 pathway.

Biological sex is an important factor that impacts both tumorigenic processes and treatment response^51–56^. Our previous studies have demonstrated that gMDSCs play a central role in driving GBM in females but did not address host factors underlying sex differences in MDSC subset activity^10^. The assessment of potential drug candidates aimed at targeting gMDSCs based on their differential expression profile identified GABA as a potential modulator. Importantly, this list also included IL-1b, G-CSF and metalloproteinase inhibitors, all of which were previously defined as gMDSC targets^14,35,36^, further reinforcing the strength of this prediction approach. Recent studies highlight the significance of metabolic reprogramming of tumor-infiltrating MDSCs for their ability to suppress immune response in GBM^12,57,58^. Moreover, GABA has been shown to regulate immune cell metabolism^31,39^. Our assessment demonstrated that GABA increased arginine metabolism specifically in female gMDSCs potentially via the observed upregulation of the arginine transporter Cat2B. Baseline L-Arginine levels and transporter expression were higher in males compared to females, providing a potential explanation for the differential susceptibility of female gMDSCs to GABA-induced reprogramming. The metabolic conversion of L-Arginine by Arg-1 and NOS2 are both pivotal for MDSC function, and NOS2 plays a critical role in fostering immune suppression^40,42,45^. In line with this, we observed a significant increase in GABBR-mediated NOS2 expression and T cell suppression by female gMDSCs. While GABA has been shown to suppress T cell activation, drive immunosuppressive macrophages and induce cancer cell proliferation, we did not observe changes in either of these readouts with GABBR stimulation^26,30,31,59^. One potential explanation for this discrepancy is the differences in GABA_A_R vs GABBR signaling in distinct cellular population and disease contexts. While these discoveries offer crucial mechanistic understandings regarding the sex-specific GABBR signaling in MDSCs, inhibition of NOS2 was not sufficient to extend survival in control mice, suggesting that NOS2 is only one of the GABBR-mediated mechanisms involved in the sex-specific GBM immune response.

Despite the success of immunotherapy in preclinical models and other solid tumors, options for treating GBM remain limited with immune checkpoint inhibitors exhibiting modest and variable success in GBM, partly due to the highly immunosuppressive microenvironment enriched in MDSCs^10,15,60,61^. Moreover, MDSCs expand in patients with GBM and correlates with worse disease prognosis^10,13–15^ highlighting the importance of targeting these cells while accounting for patient variability. Building on our observation that GABBR signaling drives GBM in females, we sought to target this pathway. Our findings revealed that blocking GABBR extended survival in female GBM in an immune-dependent manner. Reduced immune infiltration in female tumors suggest that CGP 35348 modulates tumor-specific immune responses in GBM. While further investigations are needed to elucidate the precise mechanisms underlying the therapeutic effect of CGP 35438 in GBM, we observed reduced NOS2 from female gMDSCs, highlighting its ability to reverse sex-specific pathological immune interactions. Importantly, these results provide a therapeutic opportunity to specifically target GABBR signaling to modulate anti-tumor immune response in GBM, given that GABA_A_R signaling restricts GBM cell proliferation and thus is potentially beneficial^26,29^.

These observations have additional clinical implications since GABA analogs are used for the management of seizures and neuropathic pain in cancer patients, including those with GBM^46,62^. While the precise mechanism of action and molecular targets of these drugs remain unknown, our results demonstrating that pregabalin drives female preclinical models and female GBM patients on GABA analogs have worse short-term outcomes suggest that GABA modulators may have sex-specific adverse effects on GBM. This highlights the need for future studies dedicated to understanding the impact of GABA analogs in the tumor microenvironment. In summary, our study highlights the opportunity to develop neurotransmitter modulators as potential regulators of tumor-immune interactions and underscores the importance of considering sex as a biological variable in both preclinical and clinical studies of immunotherapy.

## METHODS

### Ethical Regulations

The studies were performed in compliance with the guidelines approved by Institutional Animal Care and Use Committee (IACUC), University of Miami (Ethical approval number: 23 -057), Cleveland Clinic. Human blood was obtained from Suncoast Blood Center, Florida

### Materials

Baclofen (cat # 27236) and CGP 35348 (cat # 18599) were purchased from Cayman Chemicals. Pregabalin was obtained from Cleveland Clinic Pharmacy. Aminoguanidine (cat# 396494, γ-aminobutyric acid (GABA, cat # A2129), and Ficoll (cat # F4375) were purchased from Sigma-Aldrich. Isofluorane, Buprenorphine, Ketamine and Xylazine were obtained from the Division of Veterinary Resources (DVR) at University of Miami. RPMI 1640 with L-glutamine (Corning; cat# 45000-396), DMEM F/12 media (cat# 11320033), B-27 supplement 50X (cat# 17504044), Glutagro Supplement (Corning, cat# 24123012), and heat-inactivated fetal bovine serum (Corning; cat# 736) were purchased from VWR. IMDM media (Gibco, cat# 12440053), antibiotic-actinomycotic 100X (Gibco, cat# 15240062) and Penicillin Streptomycin 100X (Gibco, cat# 15070063) were purchased from Thermofisher Scientific. Recombinant Human EGF (cat# 585508) and recombinant Human FGF (cat# 710308) were obtained from Biolegend.

Mouse FcR blocking reagent (cat # 130-092-575), human FcR blocking reagent (cat # 130-059-091), CD8a microbeads mouse (cat # 130-117-044) were purchased from Miltenyi Biotec. CFSE cell division tracker (cat # 423801), Zombie Violet Fixable viability Kit (cat # 423114), zombie aqua (cat #,423101), true nuclear transcription factor buffer set (cat # 424401), recombinant GM-CSF (cat # 576306), and IL-4 (cat # 574306) were purchased from Biolegend while brilliant stain buffer (cat # 566385) was obtained from BD Biosciences.

In vitro mouse flow cytometry antibodies CD11b M1/70 (cat# 101212, 101206), Ly6C HK1.4 (cat# 128008, 128036), Ly6G 1A8 (cat# 127645, 1276424), NOS2 W16030C (cat# 696806), Arg-1 W210471 (cat# 165804), CD80 16-10A1 (cat# 104732), CD86 GL-1 (cat# 105040), CD163 S150491 (cat# 155308), CD204 1F8C33 (cat# 154718), and IA/IE M5/114.15.2 (cat# 107628) were obtained from Biolegend.

In vivo immune profiling mouse flow antibodies CD11b M1/70 (cat# 101243), Ly6C HK1.4 9 (cat# 128036), Ly6G 1A8 (cat# 127618), CD68 FA-11 (cat# 137026), F4/80 BM8 (cat# 123130), CD45 I3/2.3 (cat# 147706), CD11c N418 (cat# 117339), P2Ry12 S16007D (cat# 848004, IA/IE M5/114.15.2 (cat# 107628), PD-L1 10F.9G2 (cat# 124315), CD204 1F8C33 (cat# 154718), CD3 17A2 (cat# 100237), CD4 GK1.5 (cat# 100469), CD8a 53-6.7 (cat# 100750), B220 RA3-6B2 (cat# 103212), CD69 H1.2F3 (cat# 104526), CD152 UC10-4B9 (cat# 106305), PD1 29f.1A12 (cat# 135216), NK1.1 S17016D (cat# 156512), TIGIT 1G9 (cat# 142111), and NOS2 W16030C (cat# 696808) were purchased from Biolegend.

Human flow antibodies CD45 HI30 (cat# 304024), CD3 OKT3 (cat# 317336), CD19 HIB19 (cat# 302230, 302218), CD56 5.1H11 (cat# 362506, 362544), CD11c 3.9 (cat# 301636), CD123 6H6 (cat# 306006), CD11b ICRF44 (cat# 301324), Lox-1 15C4 (cat# 358606, 358610), CD14 M5E2 (cat# 301804), CD33 WM53 (cat# 303430), HLA/DR L243 (cat3 307618), CD66b G10F5 (cat# 305122), CD68 Y1/82A (cat# 333826), Arg1 14D2C43 (cat# 369708), CD4 SK3 (cat# 344646), CD8 SK1 (cat# 344704), CD45RA HI100 (cat# 304130), CD62L DREG-56 (cat# 304806), CD69 FN50 (cat# 310934), and CD95 DX2 (cat# 305646) were purchased from Biolegend. Mouse/human GABBR1 S93A-49 (cat # MA545533) and GABBR2 S81-2 (cat # MA545536) were obtained from Life Technologies. Lightning-Link Pe Cy7 antibody labeling kit (cat # 7620030) and human/ mouse/ rat iNOS antibody 2D2-B2 (cat # MAB9502) were purchased from Biotechne.

### Cell lines

SB28 line was obtained from Dr. Hideho Okada (University of California, San Francisco), KR158 and L1 cells was received by Dr. Loic Deleyrolle (University of Florida), and DI318 cells were acquired from Dr. Justin D. Lathia (Cleveland Clinic). Mouse cells were maintained in RPMI 1640 (cat # 11875119, ThermoFisher) supplemented with 10% FBS (cat # 4500-736, Corning) and 1% penicillin/streptomycin (cat # 15070063, ThermoFisher). Human cell lines were maintained in DMEM F/12 media containing 10X B27, 50ng/ml EGF, 50ng/ml FGF, 1% Glutagro and 0.5% penicillin/streptomycin. All cell lines were treated with 1:100 MycoRemoval Agent (cat # 93050034, MP Biomedicals) upon thawing and routinely tested for Mycoplasma spp. (Lonza). Cells were not used beyond passage number twenty.

### Animal Studies

Four-week-old C57BL/6 male and female mice (JAX Stock # 000664) were purchased from the Jackson Laboratory as needed and housed in either the DVR facility at University of Miami or the Cleveland Clinic Biological Research Unit (BRU) Facility. 4-week-old NOD-*Prkdc^em26Cd52^Il2rg^em26Cd22^*/NjuCrl (NCG) male and female mice (Charles River Stock # 572NCG) were purchased from Charles River Laboratories. F4-6-weeks-old B6.129P2-Nos2tm1Lau/J (NOS2 KO, JAX Stock #002609) homozygous breeding pairs were purchased from the Jackson Laboratory, and the colony was maintained in the Cleveland Clinic BRU Facility. 4-week-old NOD.Cg-PrkdcscidIl2rgtm1Wjl/SzJ (NSG) mice were bred in-house at the Cleveland Clinic.

C57BL/6 mice were intracranially injected at 4-6-weeks old with 20,000-40,000 SB28 or 10,000–25,000 KR158 cells while NCG/NSG mice were injected with 10,000 SB28 or 15,000 KR158 in 5-10 μl RPMI null media into the left cerebral hemisphere 2 mm caudal to the coronal suture, 3 mm lateral to the sagittal suture at a 90Åã angle with the murine skull to a depth of 2.5 mm, using a stereotaxis apparatus (Kopf). Concomitantly, mice at the University of DVR facilities were subcutaneously injected with 3.5mg/kg buprenorphine. Mice were monitored daily for neurological symptoms, lethargy and hunched posture that would qualify as signs of tumor burden.

Mice implanted with SB28 and KR158 were intraperitoneally injected with 25 mg/kg pregabalin, 2 mg/kg Baclofen, 4 mg/kg CGP 35348, or PBS (vehicle) for up to 2 weeks. The first dose was administered 7 days post tumor implantation for SB28 and or 14 days post tumor implantation for KR518, on 5 days on – 2 days off cycle. For combinatorial studies, 100 mg/kg aminoguanidine was administered intraperitoneally 7 days post tumor implantation for 5 days. Drug and vehicle-treated mice were co-housed to limit cage effect.

### Drug prediction

MDSC gene expression and network medicine analyses were previously performed and published^10^. Candidates were sorted based on z-score from low to high. The top 20 clinically-approved drugs predicted to selectively target gMDSCs were manually annotated and classified into subgroups based on their molecular targets.

### Reanalysis of single cell RNA-sequencing

GABA and GABA receptor expression from 8 male and 13 female patients with both low grade and high-grade gliomas were analyzed using the dataset GSE117891^38^ across all immune populations. Raw data was downloaded from Gene Expression Omnibus (GEO) under the accession number GSE117891 [https://doi.org/10.1093/nsr/nwaa099] and processed as described by [https://doi.org/10.1093/gigascience/giaa109]. The average expression of GABA-related gene signature (GABA_A_R [GABRA1-6, GABRB1-3, GABRD, GABRE, GABRG1-3, GABRP, GABRQ, GABRR1-3], GABBR [GABBR1-2], GABA transaminase [ABAT, ALDH5A1], and GABA transporter [SLC6A1/ SLC6A11-13]), GABBR signature and GABA_A_R signature were evaluated. Differences in immune cell populations, GABBR1 and GABBR2 in immune cell populations between male and female were compared using Wilcoxon Rank-Sum test. The analyses were performed using R (version 4.2.1).

### Effect of GABA analogs on GBM prognosis

A retrospective analysis of adult patients surgically treated for GBM from August 8^th^, 2011, to September 8^th^, 2023, was conducted. The inclusion criteria comprised of patients with pathologically-confirmed primary GBM Grade IV IDH-wildtype. Excluded were patients diagnosed with GBM Grade IV Glioma IDH-mutant, those who had undergone previous resection, individuals who had received prior chemoradiation, and those who had undergone procedures other than craniotomy for tumor resection. Patients were subsequently categorized into male or female groups and further divided based on their drug status: those who were currently using or had previously taken pregabalin or gabapentin, and those who had not. Statistical analysis was conducted using Python version 3.11.5 for MacOS. Twelve-month survival probabilities were estimated using the Kaplan-Meier method with log-rank tests for each distinct group. The event was death, and the timeline was overall survival. Overall survival was defined as date of last follow-up (or date of death) subtracted from date of surgery. Patients lost-to-follow up or had no date of death prior to the 12-month period (4 females on drugs, 16 control females, 1 male on drugs, and 24 control males) and all patients after the 12-month period were censored.

### Metabolomic screening

Bone marrows were flushed from the femur and tibia of 4-8-week-old male and female mice with 10 ml PBS and strained through a 40 μm strainer (cat # 10054, Corning). Bone marrow-derived cells (BMDCs) were incubated with 1:50 diluted mouse FcR blocking reagent in FACS Buffer (2% PBS-BSA) on ice for 5 min and stained with a cocktail anti-CD11b/Ly6G/Ly6C antibodies (1:100) for 15 min at 4°C to sort for mMDSCs (CD11b+Ly6C+Ly6G-) and gMDSCs (CD11b+Ly6C-Ly6G+) with BD FACS SORP Aria Fusion (BD Biosciences). Sorted cells were reconstituted in RPMI media (RPMI 1640 + 10% FBS+ 1% anti-anti) containing 50 ng/ml GM-CSF and IL-4. 1 million cells were treated with 100 μM GABA or equal volume of vehicle (PBS) for 18 hr in 6-well plates. The cells were harvested as pellets and snap frozen until further processing. Snap frozen cell pellets were lysed in methanol: water (80:20) at dry-ice temperature for metabolomics analysis. The analysis was performed in biological triplicate. The quantity of the metabolite fraction analyzed was adjusted to the cell count. Extracts were clarified by centrifugation, dried by nitrogen blower and then reconstituted in equal volumes 50:50 methanol: water. Metabolite fractions was processed and analyzed by targeted LC-MS/MS via dynamic multiple reaction monitoring (dMRM). All LC-MS/MS experiments were performed on an Agilent Technologies Triple Quad 6470 LC-MS/MS system with a 1290 Infinity II LC Flexible Pump (Quaternary Pump), 1290 Infinity II Multisampler, 1290 Infinity II Multicolumn Thermostat with 6 port valve and 6470 triple quad mass spectrometers. Agilent MassHunter Workstation Software LC/MS Data Acquisition for 6400 Series Triple Quadrupole MS with Version B.08.02 was used for compound optimization and sample data acquisition. Studies were performed in negative ion acquisition mode with ion-pairing chromatography, using an Agilent ZORBAX RRHD Extend-C18, 2.1 x 150 mm, 1.8 µm and ZORBAX Extend Fast Guards for UHPLC separation. Agilent MassHunter Workstation Quantitative Analysis for QQQ Version 10.1, Build 10.1.733.0 was used to integrate and quantitate metabolite peak areas. LC-MS peaks corresponding to metabolites with coefficients of variation greater than 0.5 underwent manual inspection and integration. The data was normalized to the average sum of metabolites from all the samples and analyzed using Morpheus to generate the heatmap. Metaboanalyst was used for pathway enrichment analysis.

### NOS2 and Arg1 expression in mouse myeloid cells

BMDCs were isolated from 4-8-week-old male and female mice, as described above. 250,000 cells were seeded in round bottom 96-well U-bottom plate (cat # 3788, Corning) in 100 μl RPMI media and treated with 100 μM GABA, 100 μM Baclofen or vehicle (PBS) for 18hrs. Post-treatment, cells were stained with Zombie Violet viability dye diluted 1:500 in PBS for 15 mins at room temperature. Cells were washed with wash buffer (PBS-2%BSA) and incubated with 1:50 mouse FcR blocking reagent for 5 mins on ice. Cells were then stained with anti-CD11b, anti-Ly6G, and anti-Ly6C (1:100 diluted in wash buffer) for 15 mins at 4°C. Following a wash step, cells were fixed using true nuclear fixation transcription factor buffer set. For intracellular staining, anti-NOS2 and anti-Arg1 were diluted 1:100 in permeabilization buffer and incubated with cells for 20 mins at room temperature. Cells were acquired using Beckman Coulter CytoFlex LX and analyzed using FlowJo (Version 10.10.0, FlowJo LLC).

### NOS2 and Arg1 expression in human myeloid cells

Blood samples from healthy donors were purchased from Suncoast Community Blood Bank Inc, Florida. Blood was diluted with PBS by plunging 30ml PBS through the leukoreduction system chambers using an 18 ½ gauge blunt needle (BD Biosciences act# 305180). Blood was diluted with PBS to a total volume of 45ml and that was then layered sequentially over Ficoll (1:1 ratio). PBMCs (buffy layer) were collected following centrifugation at 975*g* (9 acceleration, 0 brake) for 20 mins and resuspended in 50ml PBS, centrifuged at 450*g* for 7 mins. Following centrifugation, RBC lysis was performed by resuspending the PBMCs in 5 ml of 1X RBC lysis buffer (RBC lysis buffer 10X; Biolegend cat# 420301) for 5 mins at room temperature. PBMCs were the resuspended in 10 ml RPMI media and filtered through a 40 μm nylon mesh filter (Corning; cat# 451730).

For analyzing NOS2 and Arg-1 expression, 3-5 million PBMCs were seeded in a 6-well plate in 2 ml media and treated with 100 μM Baclofen or vehicle (PBS) for 18hrs. As described above, cells were stained with live/dead viability stain and incubated with 1:50 human FcR blocking reagent. Cells were then stained with 1:100 diluted human myeloid panel antibody cocktail and incubated for 15 mins at 4°C. Cells were fixed using true nuclear fixation transcription factor buffer set, as described above and cells were intracellularly stained with fluorophore-conjugated anti-CD68, anti-NOS2, and anti-Arg1 antibodies (1:100 diluted in permeabilization buffer). Cells were acquired using Beckman Coulter CytoFlex LX and analyzed using FlowJo (Version 10.10.0, FlowJo LLC).

### In vivo immune profiling

SB28-bearing male and female mice treated with 4 mg/kg CGP 35348 or vehicle as detailed above. All the animals were euthanized when the first mice exhibited endpoint symptoms. Blood was collected by cardiac puncture into EDTA-coated mini blood collection tubes (Greiner bio-one; cat# 450470) and centrifuged at 2000*g* at 4°C for 10 minutes. Cell pellets were resuspended in 200 μl PBS and transferred to a 96-well plate. Spleens were isolated and single cell suspensions were prepared by mechanical dissociation over a 40 μm strainer Cells were centrifuged at 340*g* for 5 mins and RBCs were lysed using RBC lysis buffer. Splenocytes were resuspended in 2 ml PBS and 200 μl was transferred to a 96-well plate. Tumors were collected and minced into small pieces with a razor blade and incubated with collagenase IV (cat # 7900, Stem Cell Technologies) containing DNase I (1:1000; cat # 18047019, ThermoFisher Scientific) at 37°C for 15 mins. Following incubation, cells were strained over a 40 μm strainer and centrifuged at 340*g* for 5 mins. Pellets were resuspended in 200 μl PBS.

For immune profiling, samples were centrifuged at 340*g* for 5 mins and washed with 200 μl PBS. As described above, samples were stained with 1:500 diluted Zombie Aqua for 15 minutes at room temperature. Following a wash step, cells were resuspended in 1:50 mouse FcR Blocking Reagent for 5 mins on ice and 1:100 diluted fluorophore-conjugated antibodies were added and further incubated for 15 minutes at 4°C. Samples were washed with PBS/BSA and fixed overnight in true nuclear transcription factor buffer set. Nex day, for intracellular staining, anti-NOS2 antibody was diluted at a ratio of 1:100 in permeabilization buffer and incubated at room temperature for 20 minutes. Samples were acquired with Beckman Coulter CytoFlex LX, and analyzed using FlowJo (Version 10.10.0, FlowJo LLC).

### T cell proliferation assay

T cells were isolated from splenocytes using CD8a microbeads and following the manufacturer’s instructions. CD8+ T cells were stained with 5 μmol/L carboxyfluorescein succinimidyl ester (CFSE) for 5 minutes at 37°C and washed with ice-cold 10% RPMI twice. 200,000 T cells were cocultured with at a ratio of 1:1 with mMDSCs, 4:1 or 8:1 with gMDSCs, and 1:1 with nonMDSCs in the presence of 100 IU recombinant IL-2 plus mouse anti-CD3/CD28 Dynabeads (cat # 11453D, Gibco) for 4 days in a 96-well round-bottom plate. For NOS2 inhibition, 0.1mg/ml aminoguanidine was added. 100μM baclofen was added to stimulated and MDSC-cocultured T cells every day. On day 4, samples were stained with anti-CD3. CFSE dilution was analyzed from CD3+CD11b− cells using Beckman Coulter CytoFlex LX.

### Macrophage polarization

BMDCs were isolated from 4–6-week-old mice, as described above. 250,000 cells were plated in 1 ml IMDM media with 20% FBS and 1% Penicillin Streptomycin containing 50 ng/ml recombinant mouse M-CSF (cat # 576406, Biolegend) in a 12-well plate. Day 3 post seeding, 1 ml media was replaced with fresh media. On day 6, cells were treated with 100 ng/ml LPS + 50 ng/ml IFN-γ, 100 ng/ml IL-4, 100 μM GABA, 100 μM Baclofen or vehicle (PBS) overnight. Cells were washed and stained for flow analysis as described above. Samples were acquired using Beckman Coulter CytoFlex LX and analyzed using FlowJo (version 10.10.0, FlowJo LLC).

### qPCR

BMDCs were isolated from 4-8-week-old male and female mice and sorted into mMDSCs and gMDSCs, as described above. Sorted cells were reconstituted in RPMI 1640 supplemented with 10% FBS, 1% anti-anti, 50 ng/ml GM-CSF and 50 ng/ml IL-4. 1 million cells were treated with 100μM GABA or vehicle (PBS) for 18hr in 6-well plates. RNA was isolated using the Qiagen RNeasy mini kit (cat # 74106) and the concentration was measured with a NanoDrop One (Thermo Scientific). cDNA was synthesized using qScript cDNA Supermix (cat # 101414-102, Quanta Biosciences) in a thermal cycler (Eppendorf). qPCR reactions were performed using QuantStudio 7 Pro applied biosystems and Fast SYBR-Green Mastermix (cat # 4385610, Thermo Fisher Scientific). For qPCR analysis, the threshold cycle (Ct) values for GABBR1, GABBR2, GABA_A_R2, and Cat2b, were normalized to the expression levels of Actin. The following primers (Integrated DNA Technologies) were used:

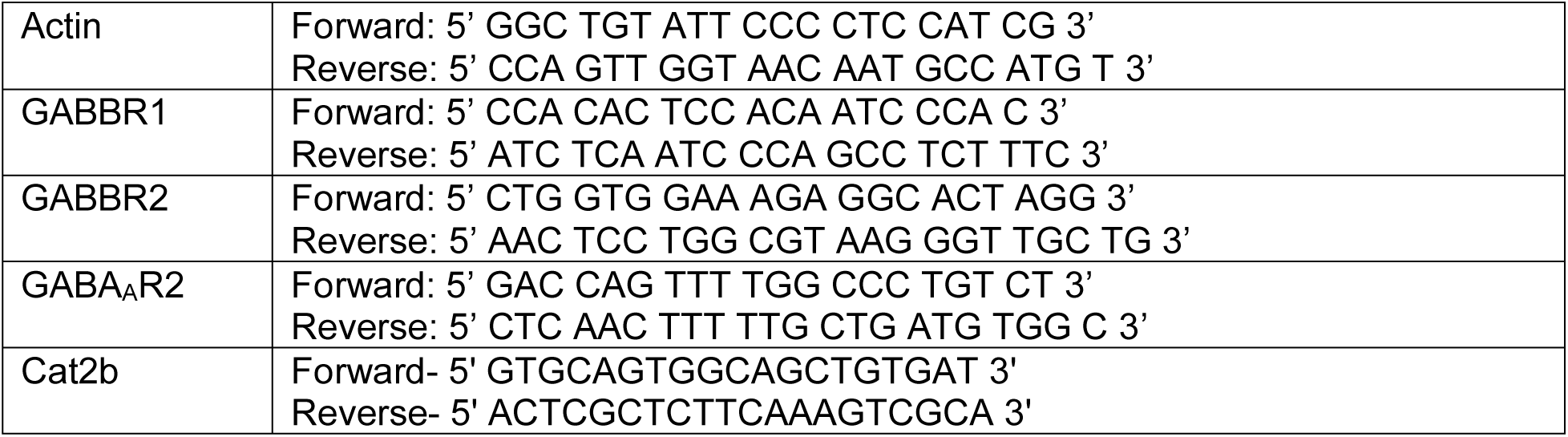

### Proliferation assay

100 SB28, KR158, L1, and DI318 cells were plated into 96-well plates coated with Geltrex (ThermoFisher Scientific) in 100 μl DMEM media supplemented with 20 ng/ml EGF, 20 ng/ml FGF, 1X B27, 1% glutagro, and 1% Pencillin Streptomycon; treated with 100 μM GABA, 100 μM Baclofen or vehicle (PBS). Cells were treated every day 5 days and luminescence measured using GloMax Discover, Promega.

### Statistical Analysis

Data was analyzed and presented using GraphPad PRISM (version 9.5.1, GraphPad Software Inc) software. Unpaired t test, paired t test, and two-way ANOVA were used for comparison of differences among sample groups. The Gehan–Breslow–Wilcoxon test was used to analyze survival data. The specific statistical method employed for individual datasets is listed in the figure legends.

## Acknowledgement

This work is supported by NCI R00 CA248611 (D.B.), American Brain Tumor Association DG 2300253 (D.B.), VeloSano Catalyst Award (D.B.), SCCC Start-up Fund (D.B., D.C.W.), American Cancer Society Postdoctoral Fellowship PF-23-1077428-01-MM (S.P.), F31 CA264849 (K.K.), NCI R00 CA277242 (D.C.W.), Forbes Institute for Cancer Discovery (D.R.W.), NCI K08CA234416 (D.R.W.), NCI P50CA269022 (D.R.W.), NCI R37CA258346 (D.R.W.), NINDS R01NS129123 (D.R.W.), the Damon Runyon Cancer Research Foundation (D.R.W.), the Sontag Foundation (D.R.W.), the Ivy Glioblastoma Foundation (D.R.W.), Alex’s Lemonade Stand Foundation (D.R.W.), the Rogel Cancer Center (D.R.W.), the ChadTough Defeat DIPG Foundation (D.R.W.), NCI P01 CA245705 (J.D.L.), NINDS R35 NS127083 (J.D.L.), the Cleveland Clinic (J.D.L.), Case Comprehensive Cancer Center (J.D.L). Research reported in this publication was performed in part at the Flow Cytometry Shared Resource (FCSR; RRID: SCR_022501) of the Sylvester Comprehensive Cancer Center at the University of Miami Miller School of Medicine, which is supported by the National Cancer Institute Cancer Center Support Grant (CCSG) P30-CA240139. We would like to thank Jack Freeman (University of Michigan, Department of Neuroscience) for their help with sample preparation. We also thank Dr. Reza Khatib for his inspiration and support.

---

**Extended Figure 1:**
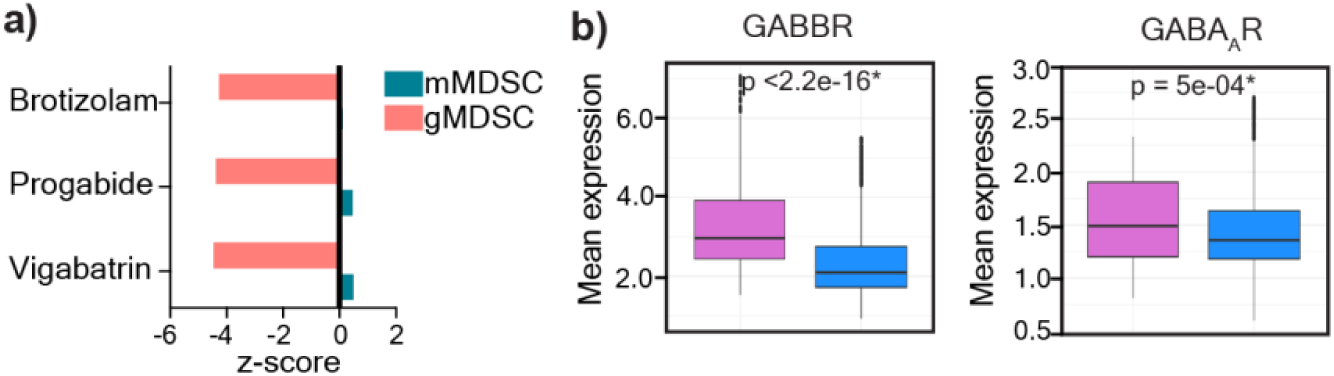
GABA receptor expression in MDSCs. **a)** Three drug candidates were predicted to target the GABA pathway in gMDSCs. Vigabatrin, a GABA transaminase inhibitor; progabide, a GABA analog, and brotizolam, a GABA receptor allosteric ligand were among the 20 candidate genes predicted to target gMDSCs based on network medicine analysis. **b)** Mean expression levels of GABBR and GABA_A_R in female (n=5) and male (n=8) glioma patients. *p<0.05 as determined by Wilcoxon Rank-Sum test.

**Extended Figure 2:**
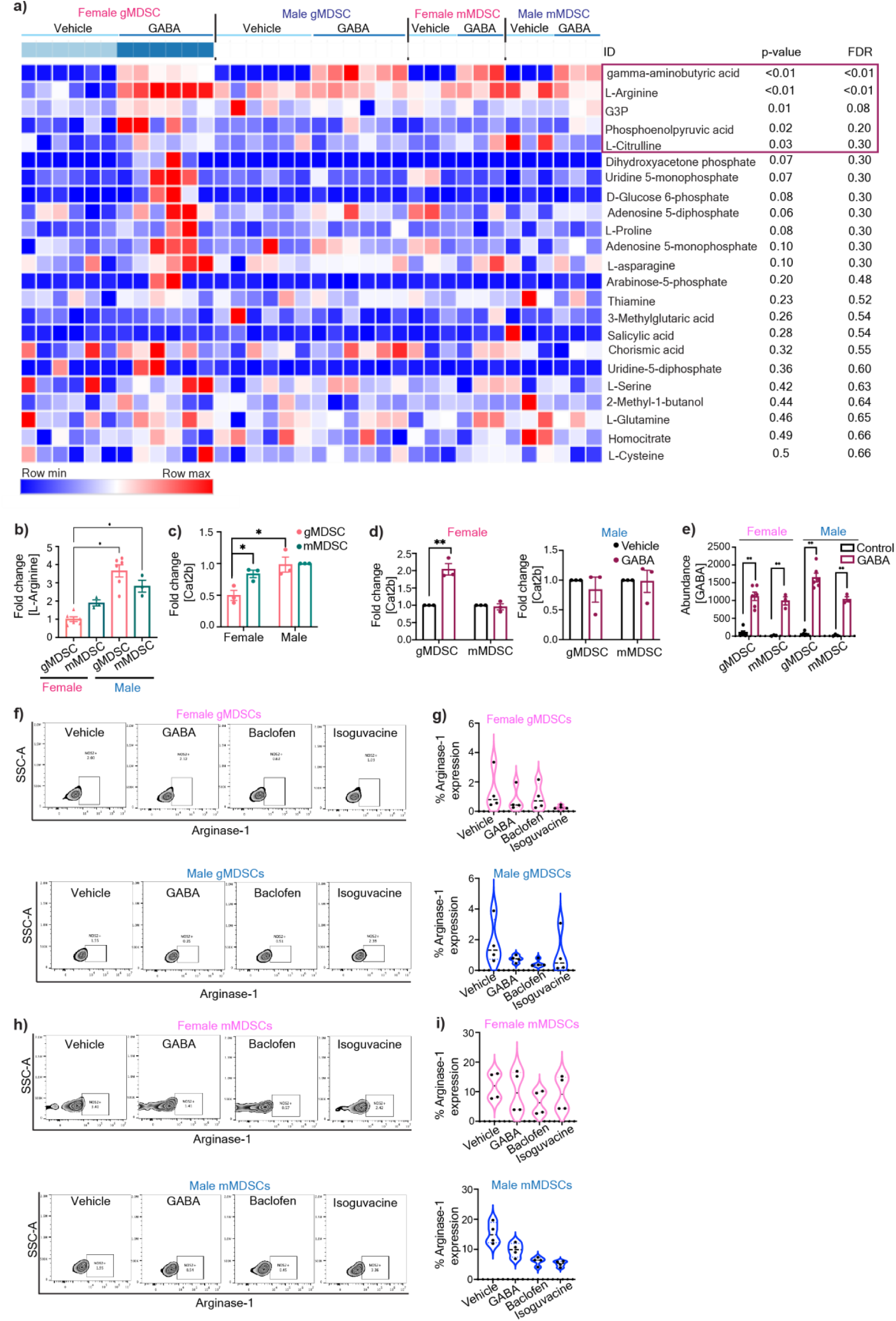
GABA-mediated metabolic alteration of female gMDSCs. **a)** Heatmap demonstrating differentially regulated metabolites in male and female mMDSCs and gMDSCs upon GABA stimulation. Data was normalized to the average sum of metabolites from all the samples and analyzed using Morpheus. **b)** Relative baseline L-Arginine levels in female and male gMDSCs (n=6) and mMDSCs (n=3), *p<0.01 as determined by two-way ANOVA. **c)** Relative expression of the L-Arginine transporter Cat2b in male and female mMDSC and gMDSC. **d**) Change in the frequency of Cat2b expression in female (left) and male (right) MDSCs treated with 100 μM GABA. n=3, * p<0.05 by unpaired t-test compared to vehicle. **e)** Enrichment plots depicting the top 25 pathways upregulated in female gMDSCs with GABA stimulation. **f)** Intracellular abundance of GABA in female and male gMDSCs (n=6) and mMDSCs (n=3) following GABA treatment, *p<0.01 as determined by two-way ANOVA.

**Extended Figure 3:**
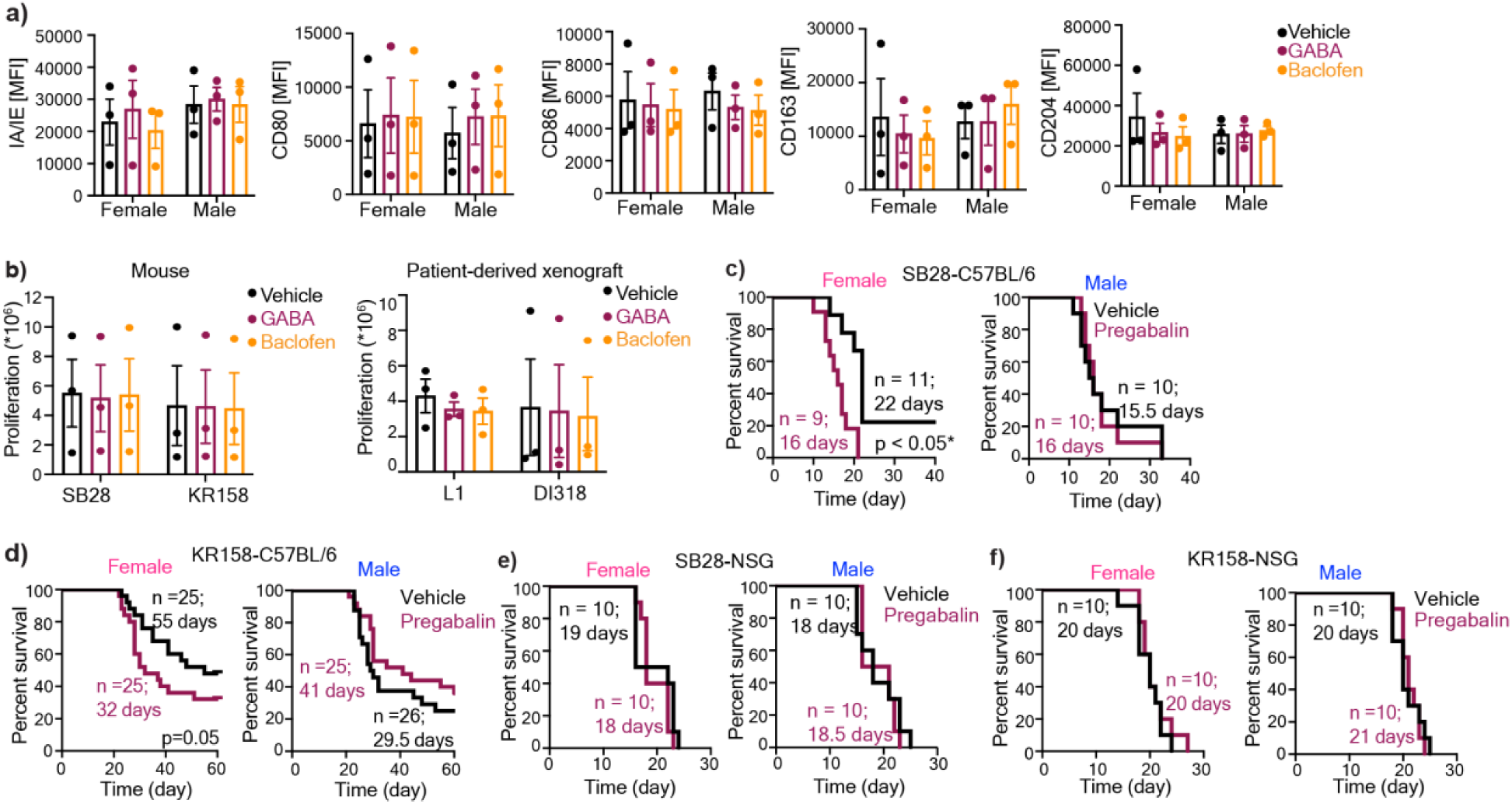
GABA analogs truncates survival in immunocompetent females. **a)** Bone marrow-derived cells were stimulated with 50ng/ml M-CSF and polarized into macrophages over a period of 6 days. Macrophages were stimulated overnight with 100 µM GABA or baclofen. Mean fluorescence intensity (MFI) of the pro-inflammatory macrophage markers IA/IE, CD80, and CD86 and the immunosuppressive markers CD163, CD204. Data shown as mean ± SEM from n=3. **b)** Proliferation rates of the mouse GBM cells SB28 and KR158 (left) and the patient-derived xenograft (PDX) lines L1 and DI318 (right), were analyzed after daily stimulation with 100 µM GABA or Baclofen. Data shown as mean ± SEM. **c)** Female and male C57BL/6 mice were intracranially implanted with 30,000 (females) and 20,000 (males) SB28 cells, or **d)** 50,000 KR158 cells. NSG mice were implanted with **e)** 10,000 SB28 or **f)** KR158 cells. Seven days post-tumor implantation, the mice were intraperitoneally injected with 25 mg/kg pregabalin daily, following a 5-days-on, 2-days-off schedule. Kaplan-Meier curves depicting the median survival of female (left) and male (right) mice. For each group Data were combined from 2-3 independent experiments. * p<0.05 based on the log-rank test.

**Extended Figure 4:**
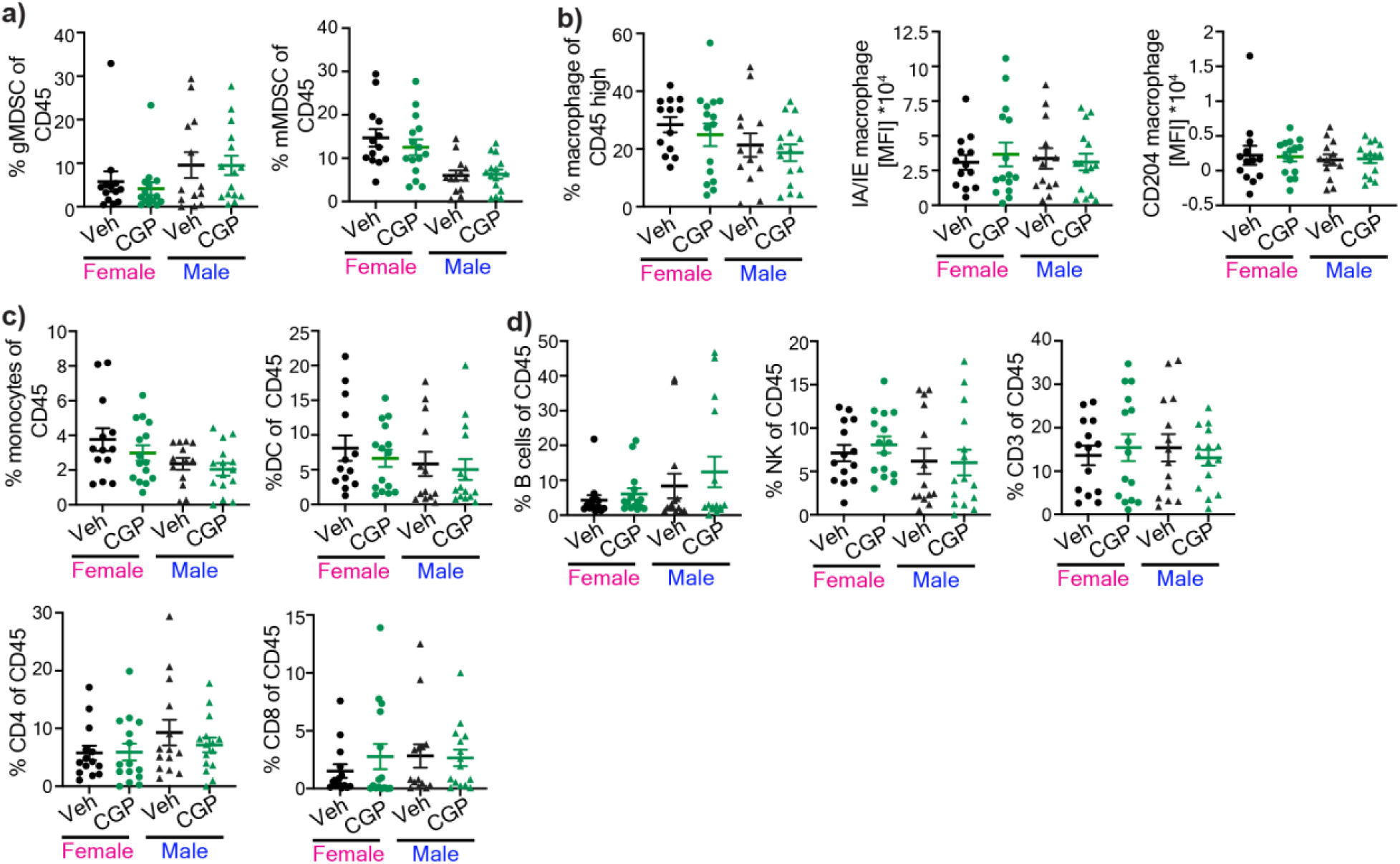
Effect of GABBR modulation on tumor-infiltrating immune populations. Tumor-infiltrating immune populations from SB28-bearing male and female mice following CGP 35348 treatment. Percentage of **a)** gMDSCs (left) and mMDSCs (right). **b**) Percentage of macrophages and MFI of IA/IE (left) and CD204 (right) of macrophages. **c)** Percentage of monocytes (left) and dendritic cells (right) **d**) B cells, NK cells, CD3+ T cells, CD4+ T cells, and CD8+ T cells (n= 13-15). Data shown from individual animals from three independent experiments.

**Extended Figure 5:**
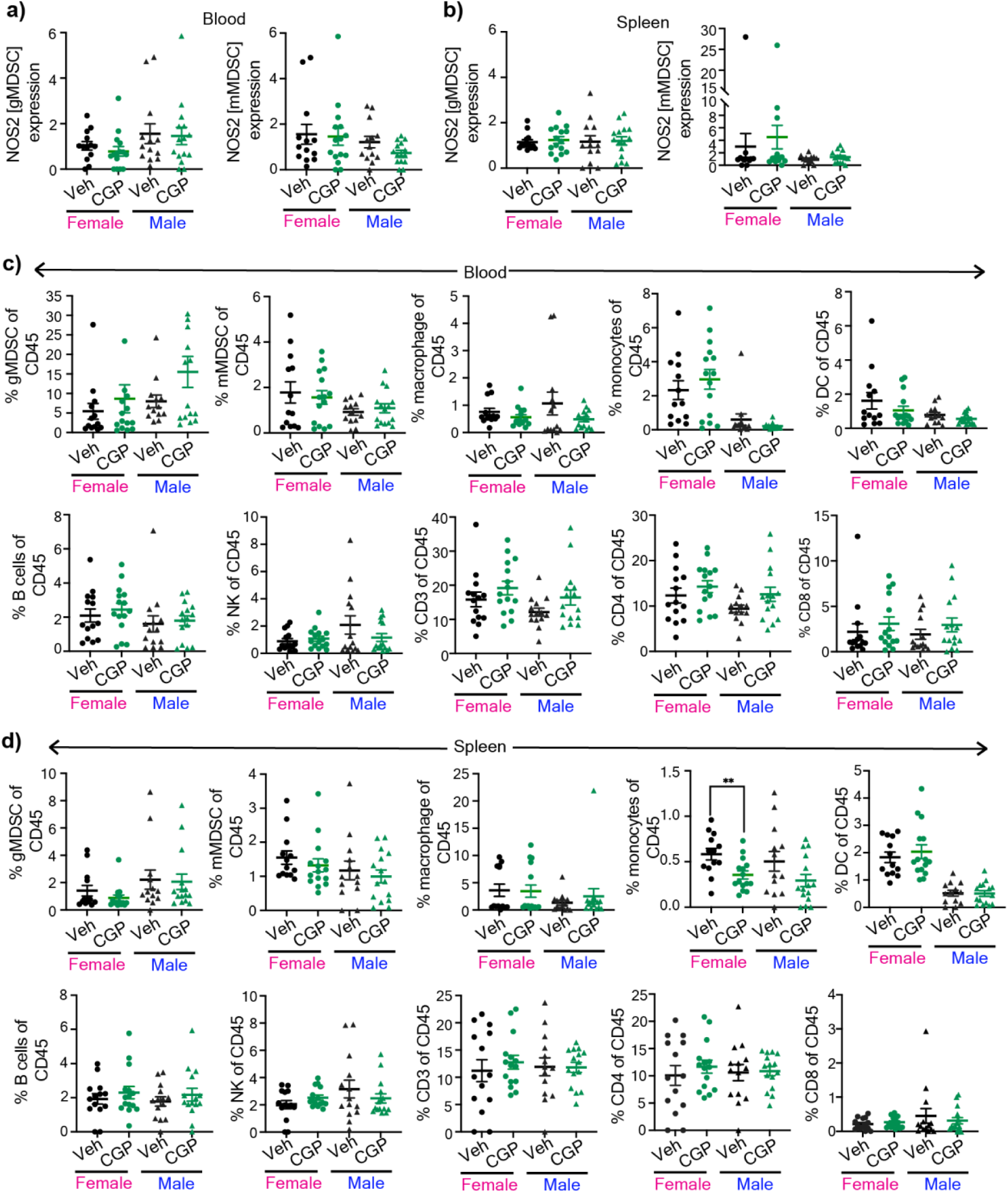
GABBR inhibition does not alter peripheral immune responses. Peripheral immune populations from SB28-bearing male and female mice following CGP 35348 treatment. Graphs showing percentage of NOS2+ gMDSCs (left) and NOS2+ mMDSCs (right) from **a)** blood and **b)** spleen. **c)** Graphs showing the percentage of gMDSCs, mMDSCs, macrophages, monocytes, dendritic cells (left to right, top panel) and B cells, NK, CD3, CD4+, and CD8+ (left to right, bottom panel) from blood. **d)** Graphs showing the percentage of gMDSCs, mMDSCs, macrophages, monocytes, dendritic cells (left to right, top panel) and B cells, NK, CD3, CD4+, and CD8+ (left to right, bottom panel) from spleen. Data shown from individual animals (n= 13-15) from three independent experiments.

**Supplementary Figure 1:**
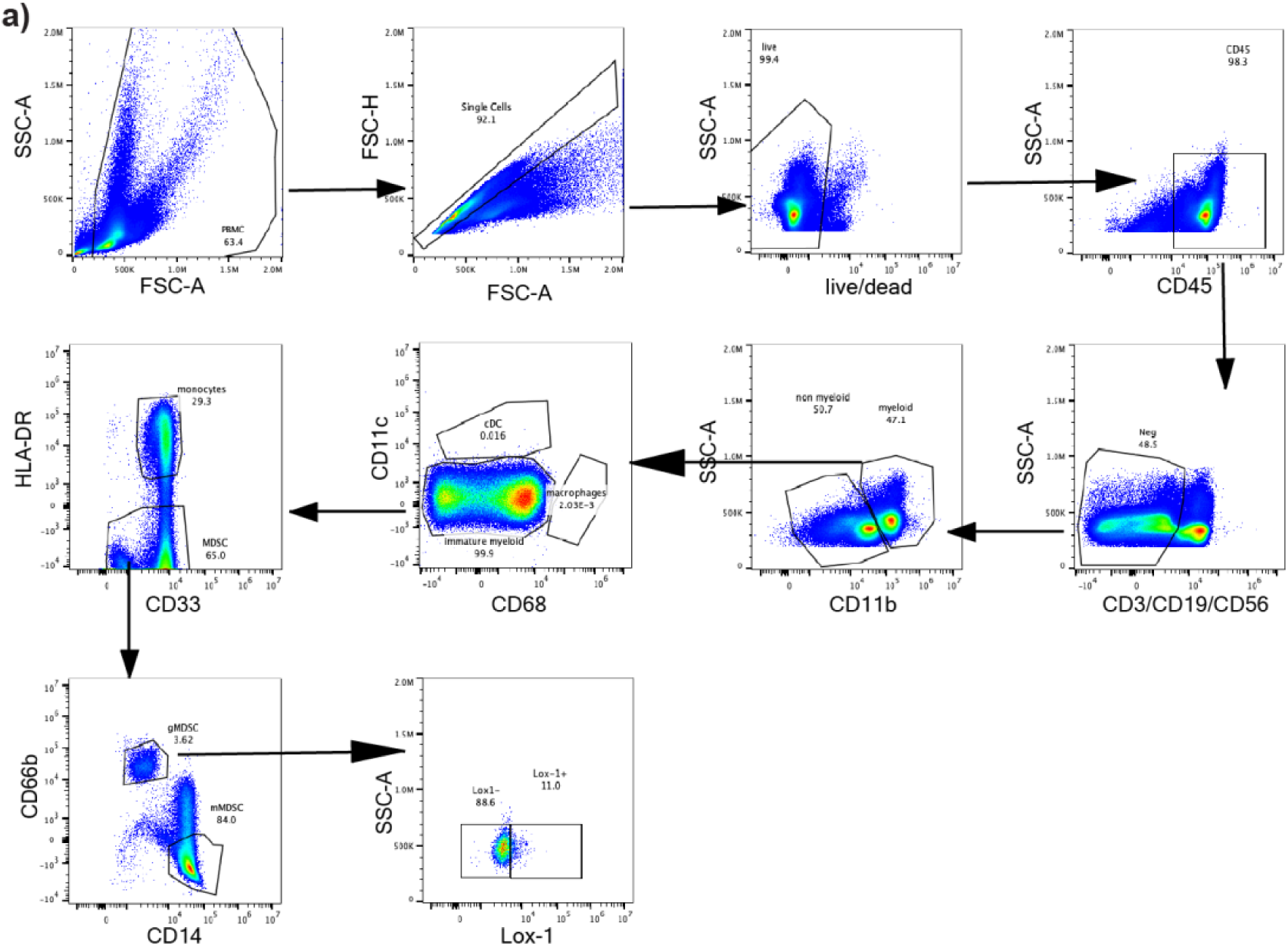
Identification of immune populations from human PBMCs. Gating strategy of immune populations from healthy human PBMCs.

**Supplementary Figure 2:**
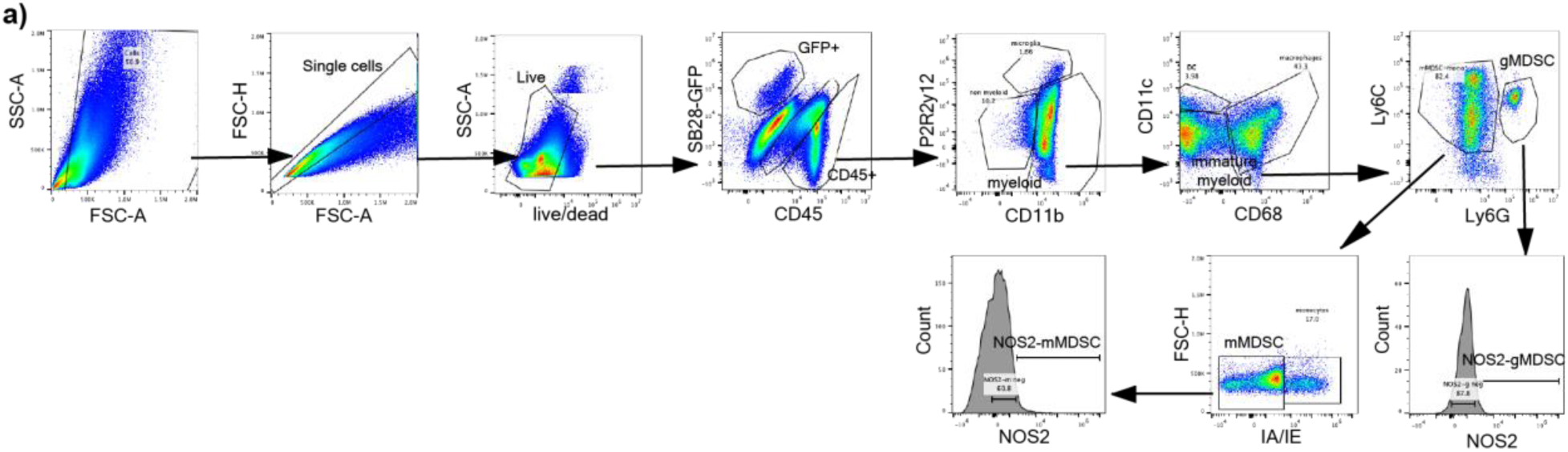
Gating strategy for immune populations from SB28-bearing mice. Gating strategy of immune populations and NOS2 expression from vehicle or CGP 35348 treated mice.

